# Imprinted loci may be more widespread in humans than previously appreciated and enable limited assignment of parental allelic transmissions in unrelated individuals

**DOI:** 10.1101/161471

**Authors:** Gabriel Cuellar Partida, Charles Laurin, Susan M. Ring, Tom R. Gaunt, Caroline L. Relton, George Davey Smith, David M. Evans

**Author notes:** **Corresponding Author:** Dr Gabriel Cuellar Partida, The University of Queensland Diamantina Institute, Brisbane, Queensland, Australia, Prof David M Evans, The University of Queensland Diamantina Institute, Brisbane, Queensland, Australia.

## Abstract

Genomic imprinting is an epigenetic mechanism leading to parent-of-origin dependent gene expression. So far, the precise number of imprinted genes in humans is uncertain. In this study, we leveraged genome-wide DNA methylation in whole blood measured longitudinally at 3 time points (birth, childhood and adolescence) and GWAS data in 740 Mother-Child duos from the Avon Longitudinal Study of Parents and Children (ALSPAC) to systematically identify imprinted loci. We reasoned that *cis*-meQTLs at genomic regions that were imprinted would show strong evidence of parent-of-origin associations with DNA methylation, enabling the detection of imprinted regions. Using this approach, we identified genome-wide significant *cis*-meQTLs that exhibited parent-of-origin effects (POEs) at 35 novel and 50 known imprinted regions (10^−10^< P <10^−300^). Among the novel loci, we observed signals near genes implicated in cardiovascular disease (*PCSK9*), and Alzheimer’s disease (*CR1*), amongst others. Most of the significant regions exhibited imprinting patterns consistent with uniparental expression, with the exception of twelve loci (including the *IGF2, IGF1R,* and *IGF2R* genes), where we observed a bipolar-dominance pattern. POEs were remarkably consistent across time points and were so strong at some loci that methylation levels enabled good discrimination of parental transmissions at these and surrounding genomic regions. The implication is that parental allelic transmissions could be modelled at many imprinted (and linked) loci and hence POEs detected in GWAS of unrelated individuals given a combination of genetic and methylation data. Our results indicate that modelling POEs on DNA methylation is effective to identify loci that may be affected by imprinting.

## Introduction

Genomic imprinting is an epigenetic mechanism in which genes are silenced in a parent-of-origin specific manner. The first experimental evidence for genomic imprinting was provided by investigations during the 1980s when researchers failed to produce viable mice embryos using only the paternal or maternal genome^1^. The precise evolutionary mechanisms that give rise to genomic imprinting are unknown. One hypothesis postulates that imprinting provides a mechanism through which maternal and paternal genomes exert counteracting growth effects during development with paternal genes encouraging growth and solicitation of maternal care, even at the expense of the mother’s health, while maternal alleles are orientated toward success of all offspring, who do not necessarily share the same father^2^. There is some empirical evidence to support of this hypothesis. For example, in contrast to expression of the paternally derived insulin-like growth factor 2 (*IGF2*) gene which promotes cell proliferation, expression of the maternally derived *CDKN1C* and *PHLDA2* genes act as negative regulators of this process^3^.

It is widely accepted that imprinted genes are regulated by *cis*-acting regulatory elements, called imprinting control elements (ICEs), which carry parental-specific epigenetic modifications such as DNA methylation^4^. DNA methylation mainly occurs at the C5 position of CpG dinucleotides and is known to influence transcription^4^. Promoter regions of imprinted genes are usually rich in CpG sites and within differentially methylated regions (DMRs); where the repressed allele is methylated and the active allele is unmethylated. Although typical imprinting of a region results in monoallelic expression of the paternal or maternal allele, studies have shown that loci can deviate from this canonical pattern and show differential expression in a parent-of-origin-dependent manner^5, 6^.

Multiple studies have shown that imprinted genes affect prenatal growth control, normal brain development and postnatal metabolism^7-10^. The monoallelic expression of imprinted loci produces genetic vulnerabilities that can lead to monogenic syndromes. In humans, abnormal imprinting patterns at specific loci can result in genetic disorders such as Beckwith-Wiedemann and Silver-Russell syndromes which primarily affect growth, and Angelman and Prader Willi syndromes which have marked effects on growth and behaviour^11^. Evidence is also growing that imprinted genes play a significant role in complex human traits. Early linkage studies found evidence that genomic imprinting was important in the genetic aetiology of mental disorders such as Alzheimer’s and schizophrenia as well as type 2 diabetes and BMI^12-14^. More recently, large-scale genome-wide association studies (GWAS) have found a handful of SNPs in imprinted genes that exhibit parent of origin effects and are associated with traits including age at menarche, breast cancer, basal cell carcinoma or type 2 diabetes ^15-18^.

Given that genomic imprinting appears to play a role in the genetic aetiology of multiple complex phenotypes, identifying novel imprinted genes is of considerable interest. However, the extent to which genes exhibit imprinted expression throughout the human genome is unknown. The number of validated imprinted genes in humans lies somewhere between 40 and 100 according to reviews^19-21^, while some databases such as geneimprint (http://www.geneimprint.com/) and the Otago imprinting database^22^ list many more that have yet to be validated. Several methods have been used to identify imprinted loci, including analysis of differential expression between parthenogenotes and androgenotes in mice^23^, bioinformatic approaches that look for novel imprinted loci based on genomic features found in known imprinted regions^24^, and creating gene knockouts of paternal/maternal alleles in mice^25^. More recently, whole genome scans of imprinted regions have been performed using next-generation sequencing technologies to measure differential gene expression between maternally and paternally derived genes using RNA-seq^26-28^ or to measure differential methylation with MethylC-Seq^29^. Although some of these more recent approaches have been applied to human genomes, the number of studies have been limited and constrained to small sample sizes^27,30,31^, thus limiting the ability to reliably detect imprinted genes.

Imprinted regions in the human genome can also be detected using statistical approaches that model parent-of-origin effects (POEs) of genetic variants on DNA methylation and gene expression. In the presence of imprinting, SNPs affecting DNA methylation (mQTLs) or gene expression (eQTLs) have a different effect depending on their parental origin. In this work, we leverage genome-wide DNA methylation and genotypic data of up to 740 Mother-Child duos from the Avon Longitudinal Study of Parents and Children (ALSPAC) to systematically identify imprinted loci. We subsequently investigated the association between SNPs in these imprinted regions and a diverse range of phenotypes.

## Methods

### Data

#### Study sample

ALSPAC is a geographically based UK cohort that recruited pregnant women residing in Avon (South West England) with an expected date of delivery between 1 April 1991 and 31 December 1992. A total of 15 247 pregnancies were enrolled, with 14 775 children born^32, 33^. Of these births, 14 701 children were alive at 12 months. Ethical approval was obtained from the ALSPAC Law and Ethics committee, and the local research ethics committees. Appropriate consent was obtained from the participants for genetic analysis. Please note that the study website contains details of all the data that are available through a fully searchable data dictionary [http://www.bris.ac.uk/alspac/researchers/data-access/data-dictionary/].

The data used in this study corresponds to the mother-child pairs from the ALSPAC cohort who took part in the Accessible Resource for Integrative Epigenomic Studies (ARIES, http://www.ariesepigenomics.org.uk/)^34, 35^. We used genotypic data from 740 mother-child duos, and DNA methylation data from the 740 children. Each child had DNA methylation measured at three time points – i.e. cord blood, peripheral blood (whole blood, buffy coats, white blood cells or blood spots) during childhood (~7 years) and during adolescence (15, 17 years).

#### DNA Methylation

Description of the DNA methylation assays can be found elsewhere^7^. In brief, genome-wide methylation was measured using the Illumina Infinium HumanMethylation450 (HM450) arrays. These arrays were scanned using Illumina iScan and the initial quality review was done in GenomeStudio. A wide range of batch variables were measured for each sample during the data generation, including quality control (QC) metrics from the standard control probes on the array. Samples failing QC were not included in the analysis. Data points with a low signal:noise ratio (detection *p* > 0.01) or with methylated or unmethylated read counts of zero were also excluded from analysis. Genotype probes in the HM450 array of the same individual at different time points were used to identify and remove sample mismatches. DNA methylation at each CpG probe was normalised using the Touleimat and Tost algorithm implemented in the R package wateRmelon^36^ to reduce the non-biological differences between probes. From the CpG probes passing QC, we selected those that have been shown to provide genuine measurements of DNA methylation as described in Naeem *et al.* ^37^, leaving 294,841 CpG probes for the analysis. Beta values (i.e. the proportion of methylation) were used for all the analyses.

#### Genotypes

Mother-child duos participating in ARIES were previously genotyped as part of a former ALSPAC study, the details of which can be found elsewhere^32,33,38^. Briefly, children were genotyped on Illumina HumanHap550 quad-chip platforms by the Wellcome Trust Sanger Institute (Cambridge, UK) and by the Laboratory Corporation of America (Burlington, USA) using support from 23andMe. Mothers were genotyped on Illumina HumanHap660W quad-chip platform by Centre National de Génotypage (Évry, FR). Standard QC was applied to SNPs and individuals. Individuals were excluded based on genotype rate (< 5%), sex mismatch, high heterozygosity, and cryptic relatedness (defined as identity by descent > 0.125). In order to remove individuals of non-European descent, principal components (PCs) were derived from LD-pruned SNPs with MAF>0.05 using plink^39^. Individuals laying 5 s.d. beyond the 1000 Genomes European population PCs 1 and 2 centroid were excluded. SNPs with a minor allele frequency (MAF) < 1%, genotyping rate <5%, or with a deviation from Hardy-Weinberg disequilibrium (p < 1 × 10^−6^) were removed from the analysis.

Genotype Imputation was performed by first phasing the genotypes using SHAPEIT V2^40^, and then imputing to the 1000 Genomes European reference panel (phase 1, version 3) using Impute (v2.2.2)^41^. Genotypes were removed if they deviated from Hardy–Weinberg equilibrium *p* < 5 × 10^−6^, MAF <5% (the high threshold was to minimize the possibility of low frequency variants producing chance parent of origin effects through statistical fluctuation) or imputation info score <0.8. Best guess genotypes were used for subsequent analyses. The final imputed dataset used for the analyses presented here contained 2,158,724 SNPs.

### Statistical analysis

#### Identifying transmission of the alleles

The crucial first step in identifying POEs is assigning alleles to their parental origin. In order to achieve this, we applied the duoHMM algorithm implemented in the software SHAPEIT V2^42^ to the most likely imputed genotypes from the ALSPAC mothers and children. This algorithm leverages linkage disequilibrium (LD) and identity-by-descent (IBD) sharing in order to phase genotypes and resolve the parental origin of alleles at each SNP. Using a custom written *Perl* script, the phased genotypes were formatted in a way such that heterozygotes where the minor allele was inherited from the mother were coded as 1, homozygotes were coded as 0 and heterozygotes where the minor allele was transmitted by the father were coded as −1. In order to confirm the accuracy of our approach to resolve the transmission of the alleles, we compared the haplotypes of the mothers and children. We observed that for each of the children, the alleles of the haplotype inferred to be the one inherited from the mother, matched to those from the mother 99.9% of the time. We attribute the 0.1% of mismatches to genotyping or imputation errors in mothers or children.

#### Regression model

In order to identify SNPs in the genome displaying POEs on DNA methylation from the three time points (birth, childhood and adolescence), we employed a regression model^6, 43^ to estimate: the additive effect *β*_A_, defined as the equal contribution of each minor allele to the phenotype; (ii) the dominance effect *β* _D_, which measures the deviation of the heterozygote from the mean phenotypic value of the two homozygotes; and the parent-of-origin effect *β* _P_, which is the mean difference between heterozygotes (i.e. the heterozygote where allele “A” is paternally transmitted, and the heterozygote where allele “A” is maternally transmitted). In matrix annotation, with intercept term *β*_0_, the mean phenotypic value for each possible genotype can be modelled as:

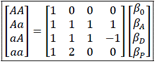

This coding of genotypes enables testing for effects that are strictly due to parent-of-origin effects, as under Hardy-Weinberg equilibrium the parent-of-origin vectors are orthogonal to the additive and dominant effects.

Given that DNA methylation is affected by sex and age, these factors were incorporated into the model as covariates, along with the first 3 ancestry informative principal components (PCs) derived from genome-wide SNP genotypes in order to control for population stratification, as well as the first 10 PCs derived from the control matrix of the Illumina HumanMethylation450 assays to control for batch effects. The following model;

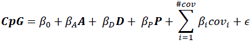

was fitted to the 294,841 DNA methylation *CpG* probes against SNPs within 500 Kb from the *CpG* probe (i.e. SNPs in *cis*). SNPs beyond 500kb from the *CpG* site were not assessed as it would have increased the multiple testing burden by 3 orders of magnitude and the number of individuals in this study may not yield enough power to detect reliable associations of SNPs in *trans*^*34*^. In this model, *CpG* is the column vector of DNA methylation values of a CpG probe; *β*_0_ is the intercept; *β*_A_ the regression coefficient of the SNP additive effect; A is the vector of genotypes in additive coding; *β*_D_ the regression coefficient of the SNP dominance effect; D is the vector of genotypes in dominance coding; *β*_p_is the regression coefficient of the SNP parent-of-origin effect; ***P*** is the vector of genotypes in parent-of-origin coding; *β*_*i*_ the regression coefficient of the covariates; and *covi* are the covariates specified above.

Given that DNA methylation values suffer from heteroscedasticity, White-Huber standard errors^44^ were computed to estimate the significance of the POE term *β*_p_ using the *sandwich* package in R.

#### Significance Threshold

In total, ~400M statistical tests were performed. Given that neighbouring SNPs usually display a high degree of correlation between each other due to LD, the number of independent tests was empirically estimated using a matrix spectral decomposition algorithm of the correlation matrix^45, 46^. We applied this algorithm in 100 randomly selected autosomal genomic regions of 1Mb each and observed that the number of independent SNPs was 0.33 times (95% CI: 0.28, 0.38) the number of SNPs tested. Hence the effective number of tests was set to 132M and the Bonferroni significance threshold was set at P-value < 3.7e-10. We note however, that this threshold may still be conservative as the correlation between CpG probes has not been taken into account.

#### Functional analyses

We performed a gene-set enrichment analysis using the WEB-based Gene SeT AnaLysis Toolkit to investigate if the identified loci implicated particular biological pathways, phenotypes and diseases. We created the gene list by mapping each CpG probe displaying a statistically significant POE to its physically closest gene. We opted for this approach over mapping genes based on whether the SNP exerting the POE on the CpG was reported to be an eQTL of a certain gene in the GTEx database, as these eQTLs are based on additive effects. Moreover, we observed that by mapping genes based on shortest distance, many of them corresponded to known imprinted genes. Gene enrichment analysis was assessed against the Gene Ontology^47^, DisGeNET^48^ and GLAD4U^49^ databases.

With a similar intent, we used Phenoscanner^50^ to identify if the SNPs exerting statistically significant POEs on CpG have been implicated in other phenotypes.

#### Predicting parental transmission in heterozygote individuals using methylation status

During this project we observed that DNA methylation at some CpG sites could potentially be used to infer the parental transmission in heterozygote individuals of samples without parental genotypes. Under a uniparental expression pattern of imprinting, one of the parental alleles remains inactive leading to the phenotypic mean of one of the heterozygote groups (e.g. minor allele inherited by the mother) being equal to the mean of the minor allele homozygote, while the phenotypic mean of the other heterozygote group (e.g. minor allele inherited by the father) is equal to the mean of the major allele homozygote. With this premise, we fitted a logistic model to the homozygous individuals for each of the statistically significant SNPs found in this study:

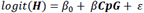

Where ***H*** is a vector with labels 0 for minor allele homozygotes and 1 for major allele homozygotes and ***CpG*** is the DNA methylation at the relevant CpG site.

We then used this fitted logistic model to predict the pattern of allelic transmission for each heterozygote individual at the putatively imprinted SNPs. Note that this approach can also predict the allelic transmission at other patterns of imprinting (e.g. bipolar or polar dominance) as it splits heterozygote individuals into those that are above the phenotypic mean of the (e.g. minor allele) homozygous individuals and those that are below the phenotypic mean of the (e.g. major allele) homozygous individuals. To measure how well this method performed, we computed the Area Under the receiver operating Characteristic curve (AUC) for each SNP.

We estimated the sample size that would be required to achieve 80% statistical power to detect POEs using this approach to infer parental transmission compared to having actual parental genotypes and being able to identify each heterozygote group correctly (as was the case in this study). We simulated 500 runs for each SNP where POEs explained: 0.5%, 1%, 2%, 4% and 9% of the variance (R^2^) using known parent-of-origin coded genotypes (i.e. 0 for homozygotes, and −1 or 1 for each of the heterozygote groups, AUC=1). We then estimated how the variance explained degraded when using the inferred genotypes coded as 0 for homozygotes and as an expected dosage for heterozygotes: *p* - (1-*p*), where *p* is the probability of being in heterozygous group 1 and 1-*p* the probability of being in heterozygous group 2. For example, when we simulated a POE using the known parent-of-origin coded genotypes (i.e. AUC=1) that explained R^2^=1%, the variance explained would drop to R^2^=0.09% when using the inferred (AUC=0.75) parent-of-origin coded genotypes (as expected, R^2^ would normally degrade relative to AUC and MAF). We then used the function pwr.r.test from the “pwr” package in R that implements a Z’ transformation of the correlation^*51*^ to derive the sample size required to achieve 80% power with α=0.0005.

## Results

### Identification of imprinted DMRs

We identified 365 CpG sites with at least one SNP exerting POEs with a p-value less than our Bonferroni significance threshold of 3.7×10^−10^ [Supplementary table 1]. These CpG sites were distributed among 85 loci, each of which was defined to be at least 2Mb apart from one another [Figure 1 and Table 1]. By inspecting RefSeq^52^, geneimprint (http://www.geneimprint.com/) and Otago imprinting^22^ (http://igc.otago.ac.nz) databases and the literature^21,53-61^, we identified 150 known imprinted loci (each defined to be at least 2Mb in each direction from one another) [Supplementary Table 2]. Of the 85 loci we identified at genome-wide significant levels, 50 mapped to these known imprinted regions, while 35 appeared to be novel. Distance between each identified locus and the closest known imprinted locus is included in Supplementary Table 3. The POEs identified were remarkably consistent across the different time points (i.e., birth, childhood and adolescence), with 70 loci identified as statistically significant at at least two time points. All the remaining loci with the exception of the *FAM30A* locus (which showed a uniparental expression pattern) showed at least a nominally significant parent of origin p-value (<0.05) between the SNP and methylation at the relevant CpG site at all 3 time points [Table 1].

**Table 1.**
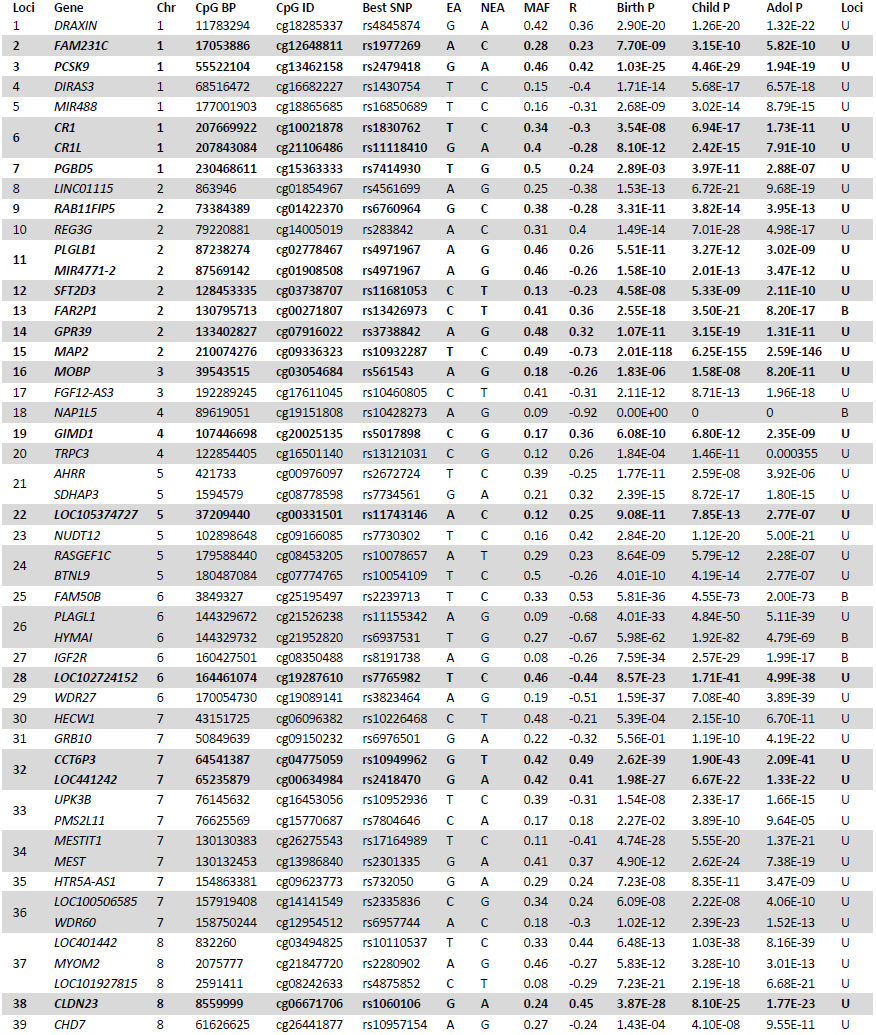
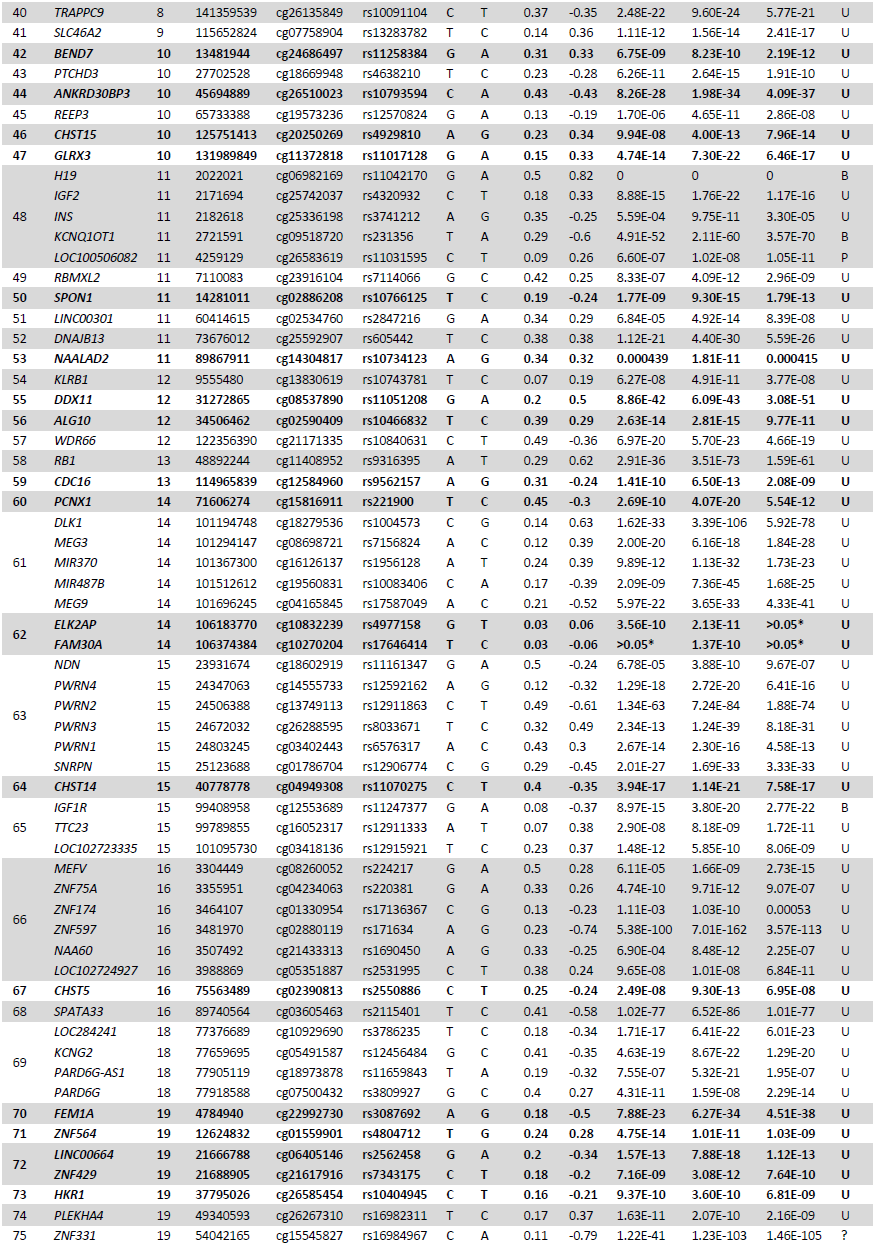
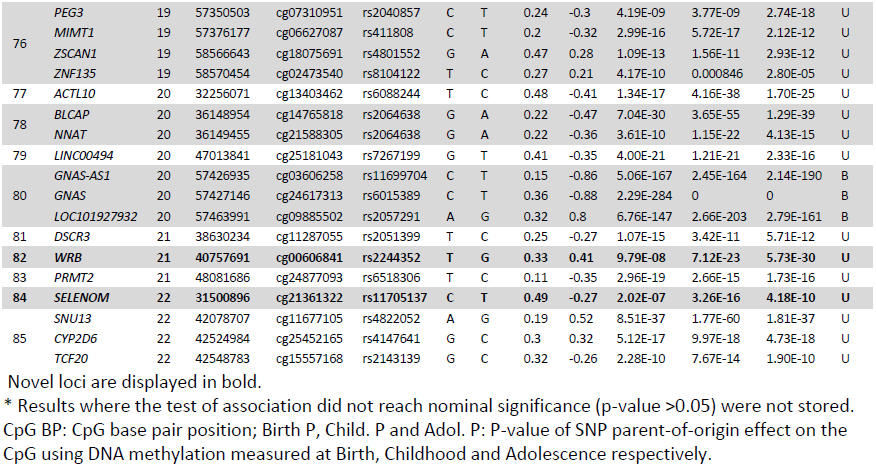
Summary of Loci identified. For each CpG site meeting experiment-wide significance, we show the SNP that produced the strongest pvalue for the POE term. If more than 1 CpG site was located at the same locus (based on distance), the one with the smallest P-value is shown. Minor alleles (MAF < 50%) were used as effect alleles (EA) while the major alleles were set to non-effect alleles (NEA). Effects are summarized as partial correlations (R) between the POE coding and methylation beta value at the CpG site. Parent-of-origin genotype coding was defined as −1 for heterozygotes where the minor allele was inherited from the father, 0 for homozygotes and 1 for heterozygotes where the minor allele was inherited from the mother. The gene reported is the one that is closest to the CpG site’s position. P-values for the POE between the CpG and the SNP are shown for each time point. Novel loci are displayed in bold. In POE patterns “U” refers to a uniparental effect, “B” refers to a bipolar pattern, “P” to a polar underdominance pattern, and “?” to an uncharacterized pattern. A definition of the different types of methylation pattern is illustrated in Figure2.

**Figure 1.**
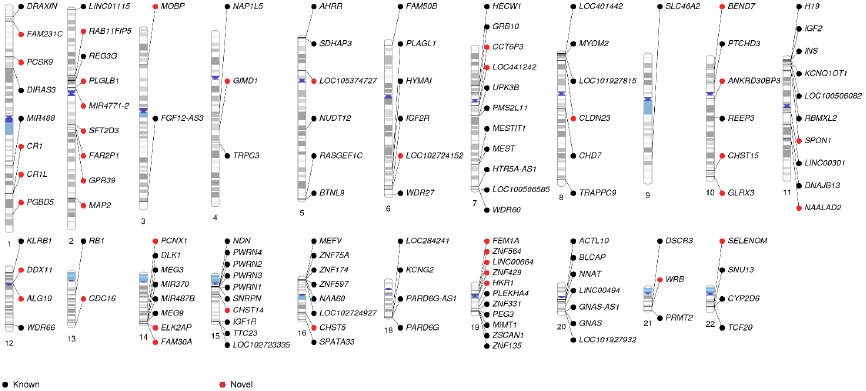
Candidate imprinted loci.

The strongest POEs were observed within known imprinted loci. For instance, we observed partial correlations (*R*) as high as 0.90 between parent-of-origin coded SNPs and CpG sites near the *NAP1L5* and *GNAS* genes. For the novel candidate imprinted loci we observed partial correlations as high as *R=0.73* for a CpG near *MAP2*. In Supplementary Tables 4-6, we have included the summary statistics of each CpG site with at least one significant SNP at each of the different time points along with additive and dominance effect statistics.

We found diverse patterns of imprinting where the effect depended on the combination of the alleles [Figure 2]. For example the distribution of DNA methylation at the CpG probe cg06982169 near the known imprinted genes *H19* and *IGF2* displayed a bipolar dominance^6^ pattern where the phenotypic value of the two homozygotes did not differ, and one of the heterozygotes had a larger phenotypic value than the two homozygotes and the other heterozygote had a smaller value [Figure 2a]. This type of pattern was also observed for loci containing the *FAR2P1, NAP1L5, FAM50B, HYMAI, IGF2R, AIRN, KCNQ1OT1, IGF1R, DDX11* and *GNAS* genes. Most of the loci identified displayed a DNA methylation distribution consistent with uniparental effects, where one of the alleles led to a larger average phenotypic value than the other and one of the chromosomes was putatively silenced. Figure 2b shows an example of this methylation pattern, where the mean DNA methylation of the CpG probe cg09336323 near *MAP2* increases only if the minor “T” is inherited from the father. We observed an instance of polar underdominance at a CpG site near *LOC100506082* where one of the heterozygous groups displayed a lower phenotypic mean than the other three genotype groups (*P-value* = 1.05E-11) [Figure 2c]. Finally, the locus containing *ZNF331* showed an uncharacterized distribution of phenotypes amongst the four possible genotypes (*P-value* = 1.46E-105) [Figure 2d].

**Figure 2.**
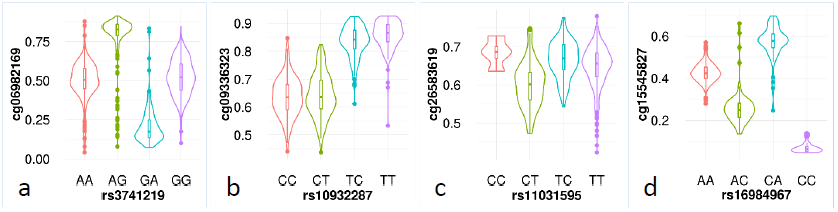
Patterns of parent-of-origin effects. Violin plots showing four patterns of CpG methylation observed in this study:(a) Bipolar dominance pattern observed at a CpG site near *H19* where one heterozygous genotype has a larger mean phenotypic value than the two homozygotes and the other heterozygote has a smaller mean value; (b) The canonical pattern of imprinting observed at a CpG site near *MAP2*, where one of the alleles leads to a larger phenotypic value than the other and one of the chromosomes is putatively silenced; (**c**) Underdominance pattern at a CpG site near *LOC100506082*, where one of the heterozygotes has a lower phenotypic value than the rest of the genotype groups; (**d**) Uncharacterized pattern at a CpG site near *ZNF331*.

### Functional analyses

To shed light on the potential function of genes within the identified loci, we performed gene enrichment analyses based on the gene ontology database^47^. Not surprisingly, we found a statistically significant overrepresentation of genes involved in insulin receptor binding (*P* = 3.63E-06; *FDR* = 6.64E-03) and imprinting (*P* = 2.76E-06; *FDR* = 2.36E-02). Beside these associations, we did not observe any other pathway with an *FDR* < 0.05; however, the top results included pathways involved in mitotic division and hormone secretion [Table 2]. Similarly, we assessed whether the genes tended to be involved in certain diseases. We found several diseases with an FDR < 0.05, most of which related to early growth or the nervous system, including neurodegenerative diseases such as Alzheimer’s and Parkinson’s disease, and syndromic disorders known to be due to imprinting aberrations, such as Beckwith–Wiedemann and Prader-Willi syndromes. These findings are summarized in Table 3.

**Table 2.** Top results of gene enrichment analysis based on gene ontology.

| GO ID | Description | Genes in set | Observed | Expected | FDR |
| --- | --- | --- | --- | --- | --- |
| GO:0005158 | Insulin receptor binding | 31 | 4 | 0.11 | 6.64E-03 |
| GO:0005159 | Insulin-like growth factor receptor binding | 15 | 3 | 0.05 | 1.54E-02 |
| GO:0071514 | Genetic imprinting | 25 | 4 | 0.1 | 2.36E-02 |
| GO:0008146 | Sulfotransferase activity | 51 | 3 | 0.17 | 4.31E-01 |
| GO:0046883 | Regulation of hormone secretion | 252 | 6 | 1.01 | 9.67E-01 |
| GO:0045840 | Positive regulation of mitotic nuclear division | 42 | 3 | 0.17 | 9.67E-01 |
| GO:0032869 | Cellular response to insulin stimulus | 188 | 5 | 0.75 | 9.67E-01 |

FDR: False Discovery Rate; Observed: Number of genes observed; Expected: Number of genes expected under the null hypothesis (no enrichment).

**Table 3.**
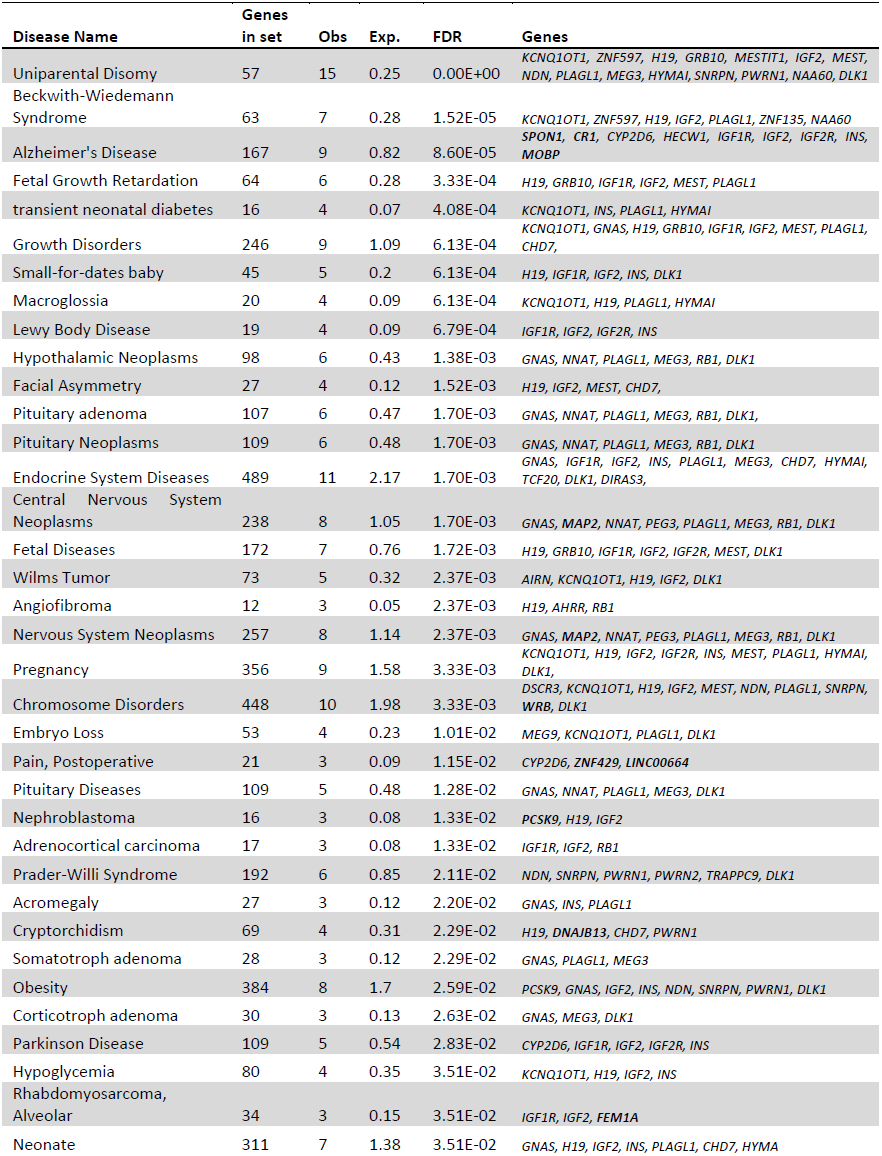
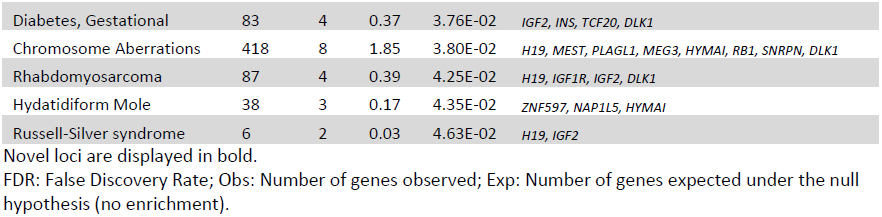
Gene enrichment analysis based ono DisGeNET and GLAD4U databases.

We also investigated whether SNPs that showed parent of origin dependent associations with methylation had also been previously associated with other phenotypes (i.e. not necessarily in a parent of origin fashion). Using Phenoscanner^50^, we identified 19 SNPs that reached genome-wide significance for POEs in our study and were also associated with one or more phenotypes (p < 5 ×10^−6^) including height, BMI, schizophrenia, fasting glucose, type 1 and type 2 diabetes **[Table 4]**.

**Table 4.** Phenoscanner lookup of significant SNPs exerting POEs.

| Phenotype | SNP | Phenotype P | CpG Chr | CpG BP | CpG ID | Birth P | Child. P | Adol. P | Closest Gene |
| --- | --- | --- | --- | --- | --- | --- | --- | --- | --- |
| Hair color red | rs258319 | 5.70E-34 | 16 | 89740564 | cg03605463 | 1.97E-76 | 4.71E-79 | 3.55E-79 | SPATA33 |
| Height | rs6088244 | 2.90E-19 | 20 | 32255988 | cg14921437 | 1.58E-19 | 4.16E-38 | 2.41E-30 | ACTL10 |
| Height | rs4320932 | 2.30E-12 | 11 | 2171694 | cg25742037 | 8.88E-15 | 1.76E-22 | 1.17E-16 | IGF2 |
| Fasting glucose | rs6976501 | 8.96E-12 | 7 | 50849723 | cg16349612 | 2.31E-05 | 1.19E-10 | 4.19E-22 | GRB10 |
| Height | rs2057291 | 2.20E-10 | 20 | 57463991 | cg09885502 | 6.8E-147 | 2.6E-203 | 2.8E-161 | LOC101927932 |
| Height | rs2735469 | 2.50E-10 | 11 | 2022021 | cg06982169 | 4.25E-31 | 1.01E-42 | 1.36E-35 | H19 |
| Type II diabetes | rs231356 | 3.70E-10 | 11 | 2721591 | cg09518720 | 4.91E-52 | 2.11E-60 | 3.57E-70 | KCNQ1OT1 |
| BMI | rs2531995 | 3.94E-10 | 16 | 3988700 | cg05351887 | 9.65E-08 | 1.01E-08 | 6.84E-11 | LOC102724927 |
| Mean corpuscular hemoglobin concentration MCHC | rs2064792 | 2.26E-09 | 6 | 164461131 | cg26079810 | 2.82E-26 | 2.57E-41 | 1.95E-38 | LOC102724152 |
| Age at menopause | rs12544305 | 8.80E-09 | 8 | 61626625 | cg26441877 | 0.000143 | 2.71E-09 | 9.55E-11 | CHD7 |
| Red blood cell traits combined | rs2238439 | 1.97E-08 | 16 | 3988869 | cg05351887 | 2.07E-09 | 3.30E-10 | 6.89E-07 | LOC102724927 |
| Serum metabolite mass spec peak 6359 mz | rs10829705 | 2.93E-08 | 10 | 131989849 | cg11372818 | 2.60E-13 | 1.52E-20 | 1.04E-19 | GLRX3 |
| Schizophrenia | rs2143139 | 4.57E-08 | 22 | 42524984 | cg25452165 | 1.99E-10 | 7.67E-14 | 1.84E-11 | CYP2D6 |
| Type 1 diabetes | rs3741206 | 6.33E-08 | 11 | 2171694 | cg25742037 | 2.47E-06 | 1.56E-11 | 3.72E-13 | IGF2 |
| Height | rs11743146 | 3.80E-07 | 5 | 37209440 | cg00331501 | 9.08E-11 | 7.85E-13 | 2.77E-07 | LOC105374727 |
| log(eGFR creatinine) | rs6593140 | 5.00E-07 | 7 | 50849723 | cg16349612 | 5.35E-18 | 5.74E-16 | 2.35E-18 | GRB10 |
| Nicotine dependence | rs171634 | 5.08E-07 | 16 | 3481970 | cg02880119 | 5.38E-100 | 7.01E-162 | 3.57E-113 | ZNF597 |
| Body mass index BMI | rs221892 | 6.96E-07 | 14 | 71606274 | cg15816911 | 1.69E-10 | 6.15E-20 | 3.60E-12 | PCNX1 |
| Spine bone mineral density BMD | rs2244352 | 9.23E-07 | 21 | 40757691 | cg00606841 | 9.79E-08 | 7.12E-23 | 5.73E-30 | WRB |
| Height | rs6976501 | 1.00E-06 | 7 | 50849639 | cg09150232 | 0.556 | 1.19E-10 | 4.19E-22 | GRB10 |
| Schizophrenia | rs4798923 | 1.07E-06 | 18 | 77659695 | cg05491587 | 9.34E-18 | 8.38E-21 | 3.61E-20 | KCNG2 |
| Height | rs2407093 | 1.30E-06 | 13 | 48892244 | cg11408952 | 3.00E-29 | 3.36E-60 | 1.32E-52 | RB1 |
| log(eGFR creatinine) | rs6593140 | 1.30E-06 | 7 | 50849639 | cg09150232 | 0.00781 | 1.59E-12 | 2.35E-18 | GRB10 |
| Hip circumference in females | rs2531995 | 1.40E-06 | 16 | 3988869 | cg05351887 | 9.65E-08 | 1.01E-08 | 6.84E-11 | LOC102724927 |
| Height | rs2531995 | 1.87E-06 | 16 | 3988869 | cg05351887 | 9.65E-08 | 1.01E-08 | 6.84E-11 | LOC102724927 |
| Serum creatinine | rs6593140 | 1.90E-06 | 7 | 50849639 | cg09150232 | 0.00781 | 1.59E-12 | 2.35E-18 | GRB10 |
| Schizophrenia | rs2240341 | 2.14E-06 | 14 | 71606274 | cg15816911 | 1.34E-11 | 6.07E-17 | 1.79E-10 | PCNX1 |
| Weight | rs2238439 | 2.22E-06 | 16 | 3988694 | cg01971227 | 1.48E-06 | 3.30E-10 | 6.22E-06 | LOC102724927 |

Phenotype P: P-value of the association between phenotype and SNP; CpG BP: CpG base pair position; Birth P, Child. P and Adol. P: P-value of SNP parent-of-origin effect on the CpG using DNA methylation measured at Birth, Childhood and Adolescence respectively.

### Using methylation to determine allelic transmissions

Given that many of the loci showing parent of origin effects were associated very strongly with patterns of methylation, we were interested in the extent to which patterns of methylation might be used to determine parental transmissions in heterozygous individuals. We examined the performance of a simple statistical approach to determining transmissions at loci showing evidence of imprinting through first modelling the methylation levels of homozygous individuals, and then using this information to estimate the transmission status of each heterozygous individual (see methods). Supplementary Table 7 displays the accuracy by which the heterozygous genotypes groups could be inferred using methylation levels at the single most strongly associated CpG site at each locus. The most accurate discrimination between heterozygote groups using DNA methylation was observed for the SNP rs11699704 near *GNAS-AS1* using DNA methylation at the CpG site cg03606258, achieving an AUC = 0.92. For the 85 loci identified in this study, the median accuracy was AUC =0.73 (interquartile range: 0.68 – 0.79).

Although for the majority of loci, the parental origin of alleles is difficult to determine with appreciable accuracy using DNA methylation alone, it may be the case that given very large numbers of individuals, it may still be possible to detect POEs in a large GWAS study of unrelated individuals when EWAS data is also present. In Supplementary Table 7 we show the sample size that would be required to achieve 80% power to detect POEs at candidate loci (α = 0.0005). The sample size required increased with lower AUC and lower MAF. For example, on average, a SNP inferred with an AUC ~ 0.75 and a MAF ~ 0.25 required a sample size 12x larger than if the SNP was inferred with perfect discrimination (AUC = 1). For more common SNPs (MAF > 0.4) and AUC ~ 0.75 the required sample would be 5x larger.

## Discussion

### Summary of imprinted loci

In this work, we presented a genome-wide scan of SNPs’ POEs on DNA methylation from peripheral blood at multiple time points. We found that most of the POEs of SNPs on DNA methylation are constant throughout birth, childhood and adolescence. This observation is consistent with previous studies, which showed that although patterns of DNA methylation at many CpG sites in peripheral blood cells are not stable over time, the additive genetic effects of SNPs on methylation appear to be remarkably consistent longitudinally^34^. We also showed that investigating POEs on DNA methylation is a powerful method of identifying regions of the genome likely to be affected by genomic imprinting. This assertion is supported by the fact that most statistically significant associations in our study corresponded to known imprinted loci and that the associations were with genetic variants in *cis* – i.e. it is unlikely that *cis* effects at genes are a product of maternal or paternal effects on children’s DNA methylation, as we expect that maternal/paternal effects to be distributed evenly over the genome and hence much more likely to be a *trans* effect than a *cis* effect.

In addition to supporting the existence of multiple imprinted genes that have yet to be characterized, we also found several instances of diverse imprinting patterns. In particular, a bipolar dominance pattern was observed among CpG sites near the insulin like growth factors and receptors *IGF1R, IGF2R* and *IGF2*, all of which are located on different chromosomes and are consistent with the hypothesis that maternal and paternal alleles are antagonistic with respect to growth^2^. Bipolar dominance patterns have been observed previously ^6, 15^ and are hypothesized to occur when differentially imprinted genes are in tight linkage disequilibrium but exert opposing effects on the phenotype **[Figure 3]**. There were also other genes nearby CpG sites that exhibited bipolar dominance POEs patterns including *GNAS* which has been previously described to encode maternal, paternal and biallelic derived proteins^62^.

**Figure 3.**
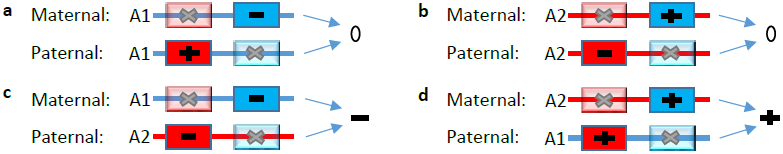
A Mechanism that generates a bipolar dominance pattern. Each of the panels in the figure displays the same two SNPs (in blue and in red) which are in high LD with each other on two different haplotypes (A1 and A2). In the case of the A1 haplotype, the allele encoded by the red SNP has a positive effect on the phenotype while the allele encoded by the blue SNP has a negative effect. In the case of the A2 haplotype, the allele encoded by the red SNP has a negative effect on the phenotype while the allele encoded by the blue SNP has a positive effect. In this example, genomic imprinting results in the red SNP being inactive in the chromosome inherited by the mother and the blue SNP being inactive in the chromosome inherited by the father. In panels (**a**) and (**b**), individuals who receive two copies of either haplotype A1 or haplotype A2 have a mean phenotype of 0. In panel (**c**) the effect on phenotype is negative as haplotype A1 is inherited from the mother and haplotype A2 from the father. In panel (**d**) the overall effect is positive as haplotype A2 is inherited from the mother and haplotype A2 from the father.

In our analyses, we identified 50 loci within the 150 known imprinted loci (summarized in Supplementary Table 2) and 35 outside these regions, deemed novel. The fact that we did not detect all known imprinted loci could be for various reasons, including lack of statistical power, poor coverage of CpG sites in the HM450 array, or the fact that imprinted expression is not maintained in all cell types^30^, and therefore we could not detect it in peripheral blood.

The strongest POEs that we identified outside known imprinted regions was on a CpG site close to the *Microtubule-Associated Protein 2* (*MAP2*) gene which plays an essential role in neurogenesis^52^. Genes located near CpGs where we also detected strong POEs included the *DEAD/H-Box Helicase 11* (*DDX11*) which is involved in rRNA transcription and plays a role in embryonic development^52, 63^, the *CCT6P4* pseudogene, and *QKI*, which is an RNA-binding protein that plays a key role in myelation^64^. Other interesting genes included *MOBP*, also involved in myelination, *CR1* which mediates cellular binding to particles and immune complexes that have activated complement, and *PCSK9*, an important gene in the metabolism of plasma cholesterol^65^.

The identification of new imprinted genes is important in furthering our understanding of the role of imprinting in normal human development and disease. The gene enrichment analyses carried out pointed to multiple metabolic, developmental and mental disorders. Alzheimer’s disease was among the top results, and was partially driven by the novel identified loci *MOBP* and *SPON1* with limited evidence in the literature to actually play a role in the disease’s aetiology^66, 67^ and *CR1*, one of the most important risk genes^68^. The SNP lookup that we performed using Phenoscanner also highlighted the potential role of the candidate imprinted loci in growth, metabolism and mental health.

### Assigning Allelic Transmissions in Unrelated Individuals

We were able to assign allelic transmissions at heterozygous individuals with moderate confidence at a handful of loci. For the remaining loci however, our predictive ability appeared to be very limited. Because of the presence of winner’s curse, these figures are likely to represent an upper limit on the predictive ability of simple approaches to resolve allelic transmission. Nevertheless, our simulations indicate that in principle POEs could be detected even if allelic transmission can’t be determined with certainty given large enough numbers of individuals with both EWAS and GWAS. Whilst there are no cohorts of this size that have this kind of information currently, it is possible that in the future as the cost of microarrays decrease, that these sorts of studies might be feasible, particularly in large scale population based cohorts like the UK Biobank where GWAS is already available^69^. Alternatively it may be possible to achieve enough power by combining cohorts with both GWAS and EWAS in a meta-analysis, as is currently being done as part of the Genetics of DNA Methylation Consortium (GoDMC). We note also that whilst we have performed power calculations using information of a single CpG per SNP, it is likely that power to detect POEs could be increased further by incorporating information from adjacent correlated CpG sites and SNPs in imperfect linkage disequilibrium.

### Strengths and Limitations

To our knowledge, this is the first study to use SNP’s POEs on human whole-genome DNA methylation to identify imprinted regions in the genome. With recent technological advances leading to decreasing sequencing costs, the current gold standard approach to identify imprinted genes is through RNA-seq – where it is possible to quantify the expression of heterozygote alleles^28,31,70^. However, this approach is still not cost effective as it is gene expression- and SNP-dependent; thus, imprinted genes with tissue-specific expression or lacking a heterozygous exonic SNP would be missed in the very small sample sizes that are common in these studies. Also, these studies usually require the genotyping or sequencing of parent-child trios in order to map the transmission of the alleles. In contrast, our approach uses large scale array data on SNPs and methylation to infer the transmission of the alleles even in absence of one parental genome. Our large sample size also provided us with greater statistical power to detect these imprinted regions.

Our approach however, does have its weaknesses. First, we were unable to directly assess whether the identified POEs directly affect the genes mentioned in this study. This is particularly problematic for the novel candidate imprinted loci where there is no prior functional work to back up our assertion. Unfortunately we could not use a more robust approach such as using eQTL information to map gene targets^71^, as eQTLs databases are based on SNPs additive effects and not POEs^72^. Therefore, the fact that we mapped genes based on their distance to the CpG sites may have impacted our gene enrichment analyses.

The other important limitation is that we were not able to distinguish whether the allele inherited from the father or the mother is active or inactive (i.e. whether the maternal or paternal gene is silenced) as the POEs are relative, and DNA methylation seldom has a baseline of zero. For instance, taking Figure 2b as an example, we cannot distinguish between whether the DNA methylation baseline is ~0.65 and the maternally inherited minor allele increases DNA methylation while the paternally derived allele remains inactive or vice versa.

The Phenoscanner lookup had the limitation that all the GWAS we looked into report additive effects, so although we observed significant associations based on additive effects, we could not assess whether these SNPs also exert POEs on these phenotypes.

### Conclusion

In conclusion, we report 35 novel genomic loci that exhibit parent of origin effects and consequently may be imprinted. We also show that the pattern of association at these loci remains stable from birth to adolescence. Although our approach does not replace traditional methods to detect genes subjected to imprinting, it is a convenient and cost-effective way to narrow down the search space and prioritize candidates. Consistent with what it is known about the biological role of imprinting, many of the identified loci were within or nearby genes with known effects on traits related to growth, development and behavior.

## Acknowledgments

This work was supported by NHMRC Project Grant (APP1085130 to D.M.E) and a Medical Research Council program grant (MC_UU_12013/1, MC_UU_12013/2, MC_UU_12013/4 and MC_UU_12013/8). The UK Medical Research Council and the Wellcome Trust (Grant refs: 102215/2/13/2) and the University of Bristol provide core support for ALSPAC. D.M.E is supported by an Australian Research Council Future Fellowship (FT130101709). Methylation data in the ALSPAC cohort were generated as part of the UK BBSRC funded (BB/I025751/1 and BB/I025263/1) Accessible Resource for Integrated Epigenomic Studies (ARIES, http://www.ariesepigenomics.org.uk). GWAS data was generated by Sample Logistics and Genotyping Facilities at the Wellcome Trust Sanger Institute and LabCorp (Laboratory Corporation of America) using support from 23andMe. We are extremely grateful to all the families who took part in this study, the midwives for their help in recruiting them, and the whole ALSPAC team, which includes interviewers, computer and laboratory technicians, clerical workers, research scientists, volunteers, managers, receptionists and nurses. This publication is the work of the authors and G.C.P will serve as guarantor for the contents of this paper.

